# Parameter Sensitivity and Experimental Validation for Fractional-Order Dynamical Modeling of Neurovascular Coupling

**DOI:** 10.1101/2021.10.20.465072

**Authors:** Fahd Alhazmi, Zehor Belkhatir, Mohamed A. Bahloul, Taous-Meriem Laleg-Kirati

## Abstract

**Goal:** Neurovascular coupling is a fundamental mechanism linking neural activity to cerebral blood flow (CBF) response. Modeling this coupling is very important to understand brain functions, yet challenging due to the complexity of the involved phenomena. One key feature that different studies have reported is the time delay that is inherently present between the neural activity and cerebral blood flow, which has been described by adding a delay parameter in standard models. An alternative approach was recently proposed where the framework of fractional-order modeling is employed to characterize the complex phenomena underlying the neurovascular. Thanks to its nonlocal property, a fractional derivative is suitable for modeling delayed and power-law phenomena.

**Methods:** In this study, we analyzed and validated an effective fractional-order for the effective modeling and characterization of the neurovascular coupling mechanism. To show the added value of the fractional order parameters of the proposed model, we perform a parameter sensitivity analysis of the fractional model compared to its integer counterpart. Moreover, the model was validated using neural activity-CBF data related to both event and block design experiments that were acquired using electrophysiology and laser Doppler flowmetry recordings, respectively.

**Results:** The validation results show the aptitude and flexibility of the fractional-order paradigm in fitting a more comprehensive range of well-shaped CBF response behaviors while maintaining a low model complexity. Comparison with the standard integer-order models shows the added value of the fractional-order parameters in capturing various key determinants of the cerebral hemodynamic response, e.g., post-stimulus undershoot.

**Conclusions:** This investigation authenticates the ability and adaptability of the fractional-order framework to characterize a wider range of well-shaped cerebral blood flow responses while preserving low model complexity through a series of unconstrained and constrained optimizations.

**Impact Statement:** The present study proposes a novel fractional-order framework for modeling neurovascular coupling. A parameter sensitivity analysis demonstrates the potential flexibility, and effectiveness of the fractional-order paradigm in reconstructing the cerebral hemodynamics with manageable complexity; and a real experimental validation analysis demonstrates the ability of the model in modeling a wider range of well-shaped CBF responses.

## I. Introduction

CHANGES in neural activity lead to changes in local cerebral blood flow (CBF) and energy metabolism. The complex relationship between neural activity and cerebral blood flow, referred to as neurovascular coupling (NVC), is a subject of intensive investigation. Understanding factors and mechanisms that orchestrate this relationship will improve our understanding of the physiological underpinnings of measurements from Functional Magnetic Resonance Imaging (fMRI) [1]. Investigating NVC in humans has become a possibility with the development of neuroimaging techniques that measure local hemodynamics, including CBF. Thus, quantitative models have been proposed to describe the different mechanisms linking transient neural activity to the changes in CBF [1]–[3]. NVC models can be arranged in a broad spectrum ranging from simple models (with fewer details and fewer parameters) to more complicated models involving many biophysical parameters and complex relationships [4]. An example of such models includes the simple model developed in Friston et al. [5], [6] and which takes a stimulus waveform and outputs a CBF response shape. The initial goal of developing this model was to fill in the neurovascular compartment to the well-known Balloon Model to predict Blood Oxygen Level Dependent (BOLD) (measured with functional Magnetic Resonance Imaging (fMRI)) responses given neural activity. A second order differential equation is used to describe CBF changes given an input stimulus. Another example is the model developed in Buxton et al. [7] which consists of a simple neural adaptation model and a linear convolution of neural activity with a flow response function.

While exploring biological processes underlying NVC using more complicated models is desired, their applicability is limited due to several mathematical constraints. For example, accurate and reliable estimation of model parameters is more difficult in complex models due to the higher number of parameters. Furthermore, parameters of those models are harder to interpret because of the lower model identifiability in complex models [4]. The difficulty of parameter estimation combined with the low identifiability and interpretability may encourage using simpler models with fewer parameters. Simpler NVC models, however, remain limited in their flexibility to fit the dynamics of CBF responses derived from specific experimental conditions or pathological populations. Deneux et al. [8], for example, showed that two of the dynamic linear NVC models (namely Friston’s model and Buxton’s model) along with their nonlinear variations were unable to capture different dynamics of CBF responses at various stimulation lengths. Linear variations appear to fail to fit the amplitude variations of different responses. While they underfit responses to short stimulations, they overfit responses to longer stimulations. Nonlinear variations also fail to account for temporal dynamics of the CBF responses especially those for shorter stimulations [8]. Those results call for developing more flexible models that can fit experimental data without compromising model simplicity.

In the last decades, non-integer differentiation, the so-called fractional-order differential calculus, became a popular tool for characterizing real-world physical systems and complex behaviors from various fields such as biology, control, electronics, and economics [9]–[11]. The long-memory and spatial dependence phenomena inherent to the fractional-order systems present unique and attractive peculiarities that raise exciting opportunities to represent complex phenomena that represent power-law behavior accurately. For instance, the power-law behavior has been demonstrated in describing human soft tissues visco-elasticity and characterizing the elastic vascular arteries. *In-vivo* and *in-vitro* experimental studies have pointed that fractional-order calculus-based approaches are more decent to precisely represent the hemodynamic; the viscoelasticity properties of soft collagenous tissues in the vascular bed; the aortic blood rate [12], [13]; red blood cell (RBC) membrane mechanical properties [14] and the heart valve cusp [?], [15]–[17] Consistent with the ability of fractional-order models to fit temporal dynamics, Belkhatir et al. in [18] have shown that the fractional-order model can yield better fit to Blood-Oxygenation-Level-Dependant (BOLD) signals measured with fMRI when compared with the original integer-order NVC model proposed by Friston et al. in [5]. Fractional calculus has been used as a powerful tool to understand better the dynamic processes that span spatiotemporal scales. Essentially, the fractional-order model is a continuous-time model with high flexibility to fit high-order dynamics and complex nonlinear phenomena. Based on this study, fractional calculus seems to be a suitable approach for NVC modeling. In fact, one of the most important properties of fractional-order derivatives is that they depend on the entire history of a function, not only the value of the function at the evaluated point. This property, called non-locality or memory effect, is relevant for modeling systems that exhibit temporal dynamics and delays such as CBF response since the model’s response at any given time depends on the whole history of the CBF response.

The presented work aims to extend the previous work [18] by studying the parameter sensitivity analysis of the fractional-order dynamical model of NVC and validated this model using both synthetic and real CBF data. The contributions of this paper can be summarised in two main parts:

- In the first part, the mathematical model is illustrated through a series of numerical simulations, which demonstrates the fractional-order model’s ability to fit a wider range of well-shaped CBF responses that cannot be captured with the standard models. In addition, using an extensive parameter sensitivity analysis, we study the effect of the fractional differentiation order and the model’s parameter on the CBF response.
- In the second part, the proposed model has been applied and validated using experimental CBF data obtained from [19]. Moreover, we evaluated the performance of the model and compared it to the integer-order model.

This paper is organized as follows. In Section II, we will recall some basic concepts from the fractional-order derivatives and the fractional-order neurovascular coupling mode.In addition, this section presents the parameter sensitivity analysis of the NVC fractional-order model along with the adopted method to fit the real CBF data. Section III discusses the obtained results and provides some future directions on the use of the model for analyzing the cerebral hemodynamic.

## II. Materials & Method

### A. Background

#### 1) Fractional-order Calculus

The concept of fractional calculus is very old and goes back to the seventeenth century. Fractional calculus is defined as a generalization of the integer-order integration and differentiation operators to the non-integer order. Because of its interesting properties of non-locality and memory, the interest on FD has grown in many fields of engineering and science. Examples of real-life applications include but not restricted to: viscoelastic, diffusive, biomedical and biological systems [20], [21], [22], [23].

The continuous fractional integro-differential operator 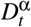, where α and *t* are the limits of the operation, is defined as follows

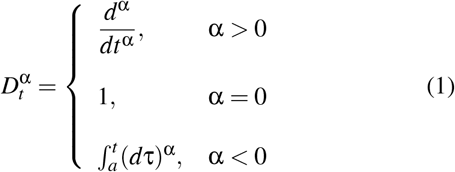

For fractional derivative, several definitions exist in the literature [24], [25]. In this work, we consider the generalized *Riemann-Liouville* (*RL*) definition which is proposed in [26] and recalled in definition 1. This definition is more appropriate for mathematical analysis.

##### Definition 1

[26] The initialized RL fractional derivative of order α ∈ (0,1) of a function *g*, denoted 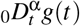, is given by:

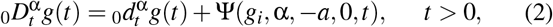

where 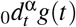 and Ψ(*g_i_*, α, −*a*, 0, *t*) are the uninitialized α*^th^* order RL derivative and the initialization function, respectively. They are given as follows:

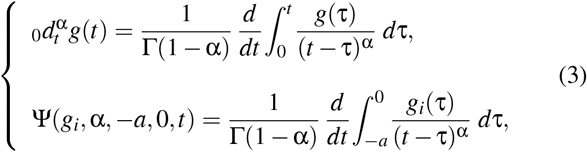

Γ(.) is the gamma function and *g_i_*(*t*) is the initialization (history) function defined for 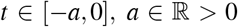. This definition assumes that the history-function for *t* ∈ (−∞, −*a*) is zero.

For numerical implementation the definition of *Grunwald-Letnikov* (*GL*) given in definition 2 is used [24].

##### Definition 2

[24] The Grunwald-Letnikov derivative of order α of a function *g*, denoted 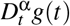, is given by:

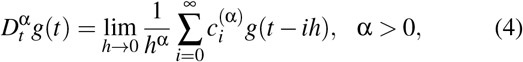

where *h* > 0 is the time step, 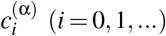 are the binomial coefficients recursively computed using the following formula,

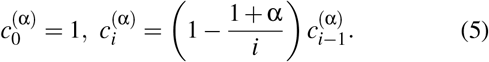

#### 2) Fractional Neurovascular Coupling Model

The proposed fractional neurovascular coupling can be formally written as:

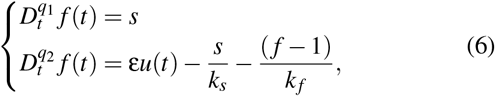

where *f* is the CBF, ε is the neural efficacy, *k_s_* is signal decay and *k_f_* is the feedback term, *q*_1_ and *q*_2_ are the fractional differentiation orders which range between 0 and 1. Note that when both fractional orders are set to 1, the model represents the original model, which we refer to as *integer-order model*, proposed by Friston et al. [5].

Fractional dynamics are only present when any fractional-order is set to a value less than 1. We refer to it as *fractional-order model*. This note is the reason why the integer-order model is a special case of the fractional-order model where both fractional parameters are set to a value of one. According to the integer-order model (where *q*_1_ = *q*_2_ = 1 in (6)), an increase in neural activity *u*(*t*) leads to an increase in the flow-inducing signal *s* (which is assumed to control CBF, *f*, at the arteriole level). The flow inducing signal *s*, then, leads to an increase in CBF, *f*. This integer-order NVC model is simple. However, and as have been pointed out in the introduction, it fails to account for temporal dynamics that arise from fractional relationships underlying CBF response [8]. Hence, two new fractional differentiation order (namely, *q*_1_ and *q*_2_) are introduced to fully model and account for the fractional properties of the CBF responses.

### B. Characterizing the unique contribution of the fractional parameters

In the first analysis, we ask whether the fractional-order model can generate unique well-shaped CBF responses that cannot be produced with integer-order models, how those contributions change the shape of the CBF response. More formally, this analysis aims mainly to exclude any equivalence that may exist between the integer-order model and the fractional-order model. In other words, we ask whether we can match any output of the fractional-order model (that has two more parameters) by solely tuning parameters of the integer-order model? If the two models are equivalent, then there exists a set of values for *k_f_* and *k_s_* in the integer-order model that can match any output of the fractional-order model. We can directly investigate this claim by running an optimization problem that minimizes signal dissimilarity between the two models. The minimization only optimizes the parameters *k_f_* and *k_s_* of both models, by fixing *q*_1_ and *q*_2_ to constant values lying between 0 and 1.

By fixing *q*_1_ and *q*_2_ of fractional differentiation order model at some constant(s) ranging between 0 and 1, we can compare the obtained optimization results (i.e. the value of the function at optimal values) across the range of *q*_1_ and *q*_2_. If the two models are essentially equivalent, then signal dissimilarity should not change across all values used for *q*_1_ and *q*_2_. However, a change in signal dissimilarity across the range of *q*_1_ and *q*_2_ indicates non-equivalence.

The cost function we used to calculate signal dissimilarity is the *L*_1_-norm of the difference between the two signals. It can be formally described as follows:

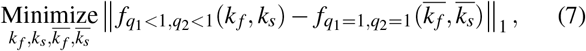

where *f*_*q*_1_<1,*q*_2_<1_ denotes the CBF computed using the fractional-order model and *f*_*q*_1_=1,*q*_2_=1_ denotes the CBF computed using the integer-order model. Each model has a separate set of four parameters. The fractional-order model has: *q*_1_ (where *q*_1_ <1), *q*_2_ (where *q*_2_ <1), *k_f_* and *k_s_*. Similarly, the integer-order model has: *q*_1_ (where *q*_1_ = 1), *q*_2_ (where *q*_2_ = 1), 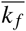 and 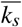. All *k_f_*, *k_s_*, 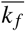 and 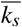 are free to vary while *q*_1_ and *q*_2_ are fixed at some predetermined values. For the integer-order model, *q*_1_ and *q*_2_ are always fixed at a value of 1 as have been noted before. However, in the fractional-order model, we use different combinations of values to test for the contribution of fractional differentiation parameters. The goal, then, is to find some combination of the four parameters that best minimize their dissimilarity, under some values for *q*_1_ and *q*_2_ in the fractional-order model.

We carried out two series of optimizations. The first one was assumption- and bounds-free where we set the lower bounds of the four parameters (i.e., *k_f_*, *k_s_*, 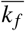 and 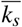) to zero and the upper bound to infinity. The aim was to numerically prove that the fractional-order model can fit CBF flow response regardless the values (or the upper bound) of the four parameters. As this leads to unrealistic CBF responses, we defined permissible ranges for the four parameters and re-ran a constrained optimization. The selection of this range (or the upper bound) followed a visual inspection of the CBF response shapes using different values for the upper bound (e.g. 100, 50, 10 and 2). More importantly, it was also guided by previous literature on the upper limits of *k_f_* and *k_s_* (i.e. the range values in [8], [27]).

### C. Parameters Sensitivity analysis of NVC fractional-order model

After showing that the fractional-order model gives raise to unique contributions to CBF response, we conduct a sensitivity analysis to study how the parameters of the fractional-order model affect or control the CBF response above and beyond the key parameters in integer-order model. To this end, we followed the sensitivity analysis approach conducted in [28]. Using a CBF response with a reference signal of fixed parameter values (shown in Table I), we slowly manipulate each parameter (and pairs of parameters) to quantify the deviation in output behavior by computing the *L*_1_-norm of the difference between the reference CBF and the manipulated CBF. Then we normalize the result by the norm of the reference signal to give more interpretable values.

**Table I:**
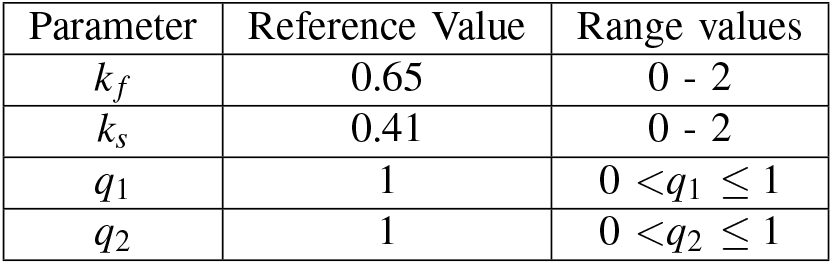
Parameters values

### D. Fitting fractional-order model to experimental CBF data

In this section, we fit the proposed fractional-order model to real CBF data obtained from another study [19]. The inputoutput data used in this paper were acquired using electrophysiology and laser Doppler flowmetry (LDF) recordings, respectively. The data acquisition of the CSD data and CBF data are described briefly in the following and for more details we refer the reader to [19], [29], [30].

Hooded Lister rats with weight’s range 200 – 300*g* were used. The animals were prepared in a way to meet certain predefined specifications. After locating the whisker barrel cortex region, electrophysiology and LDF probes were placed based on the alignment of optical imaging maps with images of the cortical surface. The inserted electrophysiology electrodes were coupled to a data acquisition device (Medusa Bioamp, TDT, FL) with a custom written Matlab interface. Field potential recording was sampled at 6103.5 Hz with 16-bit resolution. To avoid the intrinsic spatial ambiguity which is inherent in the electrophysiology data, current source density (CSD) is used [29], [31]. The CSD profiles were given to us with a sampling time between each CSD point of 200*ms*. To allow concurrent measurements of CBF, the LDF probe was positioned over the active area, adjacent to the electrodes. An LDF spectrometer including a low-pass filter was used to analyze the signal from the LDF probe with minimized errors due to the measurements noise. The CBF changes recorded with LPF were normalized to the baseline CBF which is collected for 8 *s* period before the onset of each trial. The CBF data acquired each 33.33 *ms* which correspond to an LDF with sampling rate of 30*Hz*.

Regarding the experimental paradigm, it consists of conditioning block of stimulation followed by a probing block of stimulation per trial. For each trial, two blocks of stimuli were used. The first conditioning block has three different durations (2, 8, 16 *s*) which are followed by the probing block of 1 *s* duration. These two blocks are separated by 7 different time gaps (0.6, 1, 2, 3, 4, 6, 8 *s*). Therefore, there were 21 types of stimuli paradigms run for each animal. Last, the data were animal averaged. The averaged CSD and CBF data recorded for the 21 types of paradigms are shown in Figure 4 (blue lines).

Using the CSD data as input to the model and the CBF data as its output, we evaluate how the model will fit the real-data over a certain range of parameters. To get those fittings, we first resampled the CSD input time points to the rate of CBF output data. Then, we used optimization function ’patternsearch’ (in Matlab) to find the best set of parameters that can minimize the *L*_1_ norm difference between the model outputs and the real CBF data, given the same input. The procedure was repeated for each subject, in each condition x stimulation combination. We then visualized the average fit for all subjects (while showing the standard deviation as a grey area), in each stimulation x gap combination.

## III. Results

The effects of how parameter change the CBF response have already been discussed elsewhere (see [5] for *k_s_* and *k_f_*, [18] for fractional orders *q*_1_ and *q*_2_]. Figure 1 recalls the different shapes of CBF response in a range of values for each parameter. As shown in Figure 1A and 1B, decreasing *q*_1_ reduces signal width whereas decreasing *q*_2_ increases the signal width and hence slows down signal decay. In most cases, decreasing *q*_2_ also maintains a longer post-stimulus undershoot (associated with the slower decay) as well as delayed negative peak. On the other hand, *q*_1_ only exhibits the negative undershoot in the first few fractions (approximately up to *q*_1_ = 0.8) after which CBF response returns to baseline rapidly with no apparent negative undershoot. In general, while *q*_1_ has more control over the positive aspects of the signal (i.e. overshoot width), *q*_2_ has more control over the negative aspects of the signal like signal decay and undershoot duration. Positive signal amplitude is a common feature that both *q*_1_ and *q*_2_ effectively can change. Figures 1C and 1D shows the effect of *k_s_* and *k_f_*, holding both fractional orders *q*_1_ and *q*_2_ at 1 (hence, it represents the integer-order model). Decreasing *k_s_* increases signal oscillations while increasing *k_f_* eliminates the CBF undershoot.

**Figure 1:**
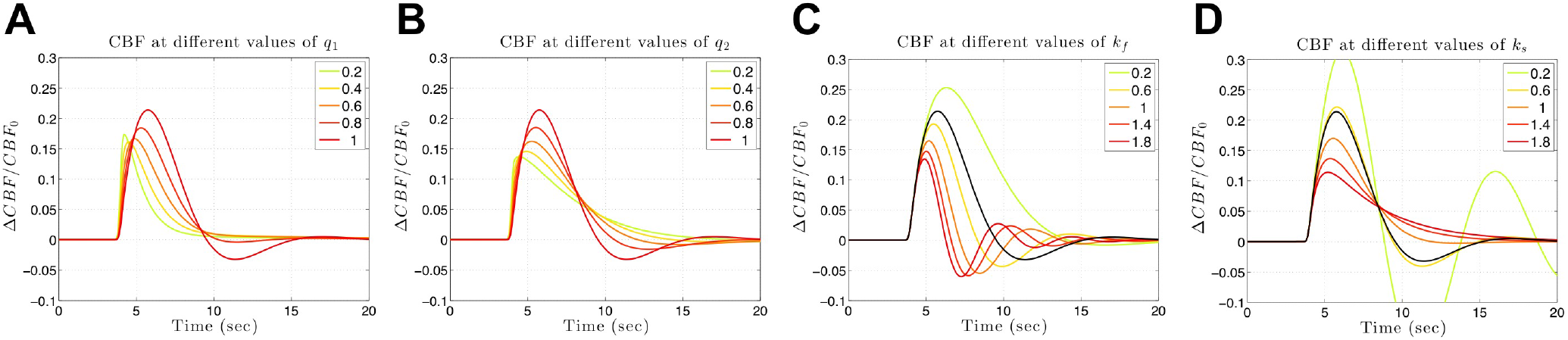
CBF response while varying each parameter. Panel A shows CBF response as a function of *q*_1_. Panel B shows CBF response as a funciton of *q*_2_. Panel C shows CBF response as a function of *k_f_*. Panel D shows CBF response as a function of *k_s_*

### A. Sensitivity analysis for the parameters of the fractional-order model

### B. Characterizing the unique contribution of fractional parameters

Figure 2 shows the results of the two series of optimizations described in Methods section. More specifically, signal dissimilarity increases as either *q*_1_ and *q*_2_ decreases. If the two models are equivalent, then a perfect match would be obtained resulting in an error of zero. However, increasing dissimilarity as a function of *q*_1_ and *q*_2_ indicates the noticeable effect of the new fractional parameters on the CBF response that cannot be obtained by tuning 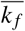 and 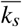 of the integer-order model.

**Figure 2:**
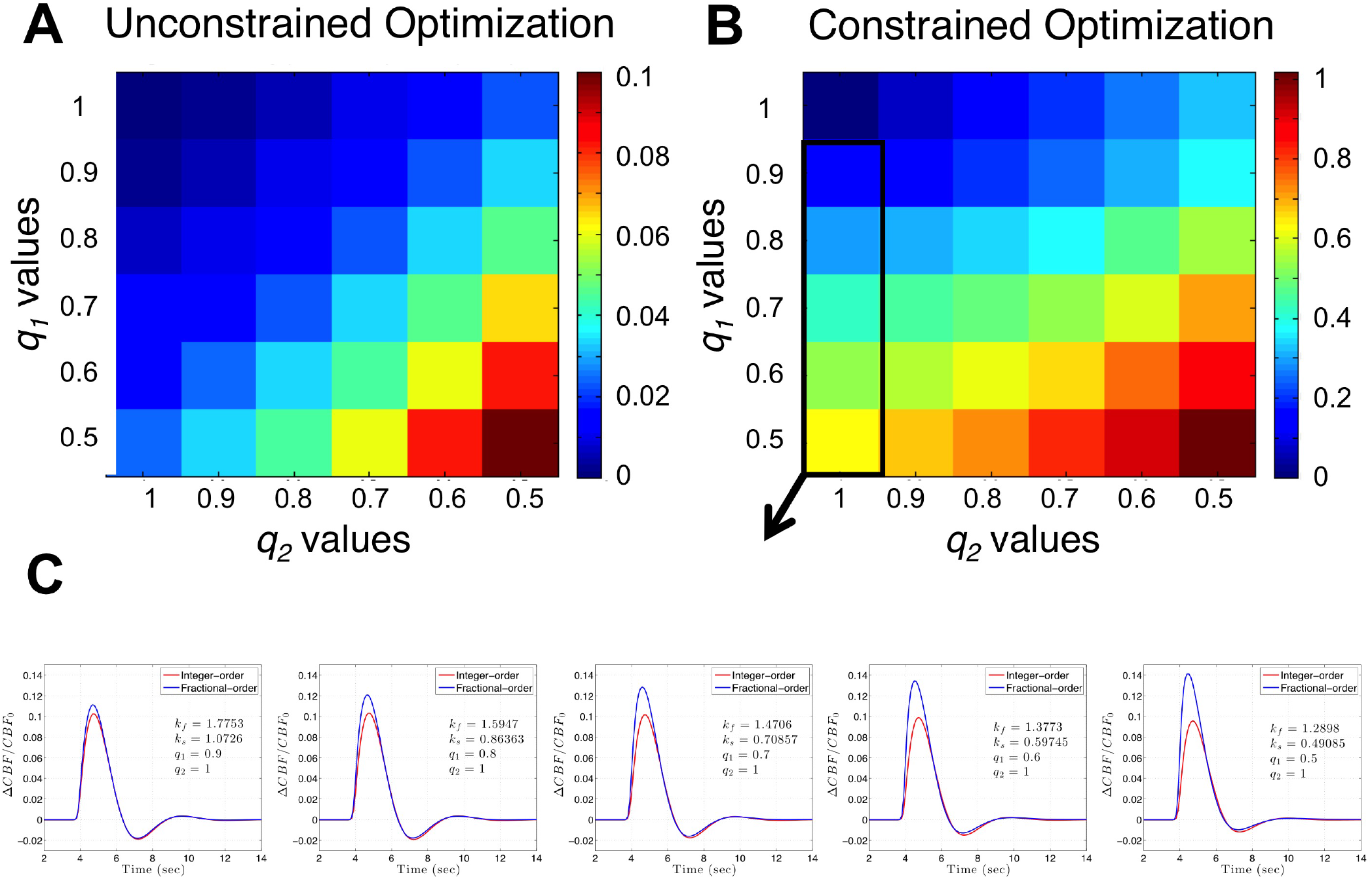
Results of the (A) unconstrained and (B) constrained optimization over *k_f_* and *k_s_* of the two models. Colors represent the L1-norm of the difference between the two CBF outputs at the optimal values. In the constrained optimization, we used range from zero to 2. Signals of both models within the black rectangle are visualized in (C)

More specifically, as we can see in Figure 2C, when varying *q*_1_ while keeping *q*_2_ at 1, we can see that CBF undershoots still matched, but overshoot amplitude is increasing due to the effect of the fractional parameter *q*_1_. We note that the actual values of error (or dissimilarity between the two models) are not of interest, but rather how they change as a function of fractional parameters.

Each subplot of Figure 3 corresponds to the relative error of the CBF signal when one (or two) of the parameters takes values in a grid around the reference value within a specific range (shown in Table I). A unique extreme in the neighborhood of the reference value of the parameter is observed for both *k_s_* (Figure 3A) and *k_f_* (Figure 3B). Notably, the relative error is asymmetric around the observed global minimum. This asymmetry indicates the CBF response becomes less sensitive to changes in those parameters as the value of either parameter increases above the reference value (see Figure 3A and 3B). Hence, initial guesses should always be taken less than the expected values of those two parameters [28].

**Figure 3:**
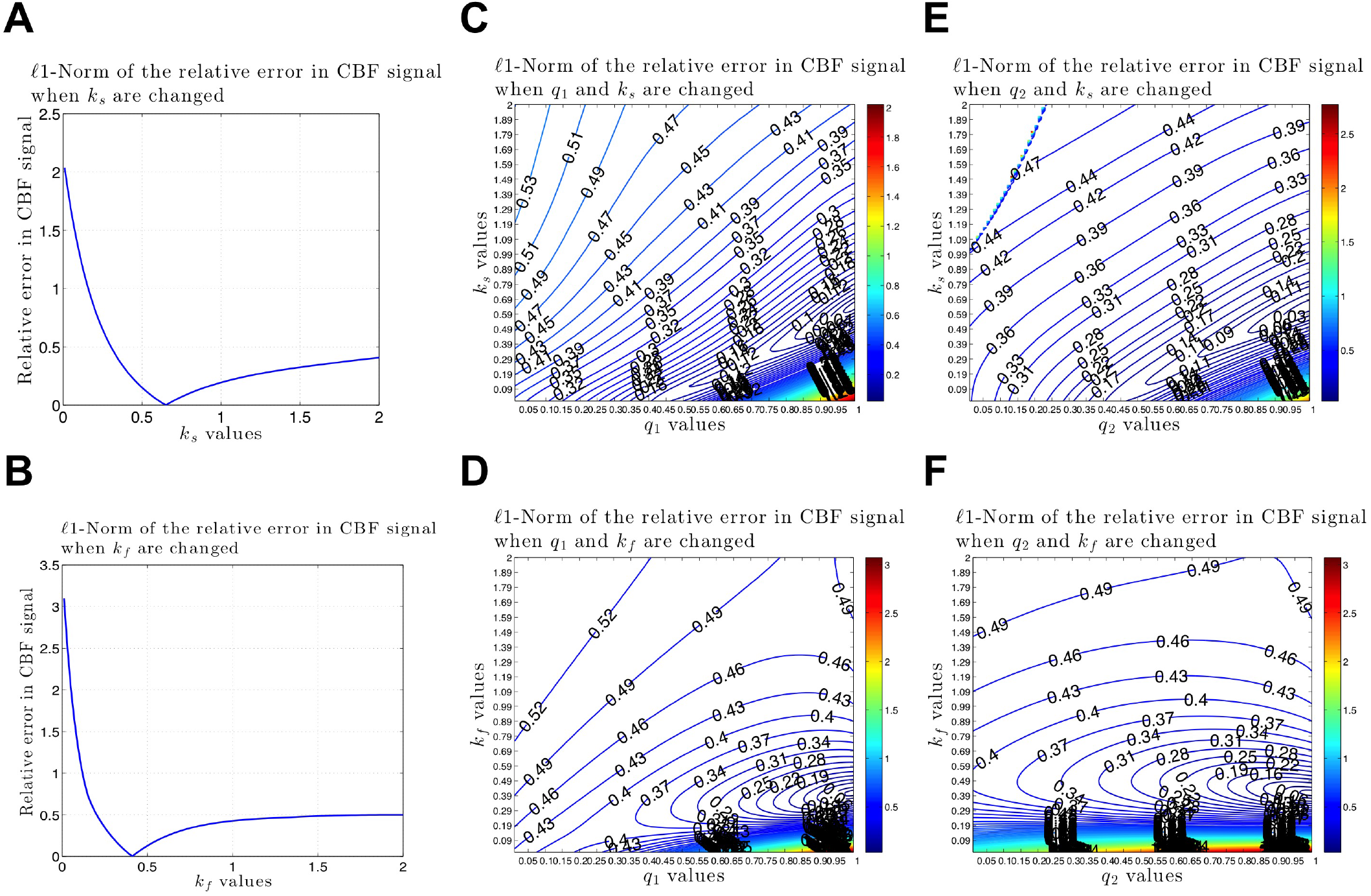
Sensitivity analysis results showing the relative CBF norm error as a function of variations in (A) *k_s_* (B) *k_f_* (C) *k_s_* & *q*_1_ (D) *k_f_* & *q*_1_ (E) *k_s_* & *q*_2_ and (F) *k_f_* & *q*_2_.

Similarly, Figure 3C - 3F shows the relative error in CBF in the case of varying two parameters (again, while keeping other parameters at their reference values). Although a global minimum is shown around the reference values, those figures show a correlation between *k_s_* and both fractional orders which indicate that *k_s_* can, to some extent, ”undo” the effects of both fractional orders. Concerning estimating the fractional parameters in light of those results, it can be argued that estimation of both *k_s_* and *k_f_* should always start with values less than their expected values as model dynamics are slower (less sensitivity) to big variations in values greater than the nominal values. For *q*_1_ and *q*_2_, initial guesses are best set to 1 as CBF is less sensitive to variations in fractional orders when they approach zero.

### C. Fitting fractional-order model to experimental data

Both integer-order and fractional-order model have been fitted to real CBF experimental data. Paired t-test of errors for each subject x condition combination show a significant difference between the errors derived from the two models (t(230) = - 5.68, p-value < 1e-7; *Mean_fractional_* = 60.706 < *Mean_integer_* = 62.913). Figures 4 illustrate the results of fitting the fractional model to experimental CBF data. Each subplot contains a combination of stimulation x gap parameters that were used in the experiment. Model fits (in blue) show a very good fit for short stimulation paradigm (2s) and a moderate fit for the 8s stimulation paradigm. However, it clearly fail to fit the long (16s) stimulation paradigm. Specifically, the model is only able to fit the second peak while missing other dynamics involved. Taken together, those results suggest that the fractional-order model has a moderately higher flexibility in fitting experimental data when compared with integer-order model due to the fact it generalizes the integer-order model.

**Figure 4:**
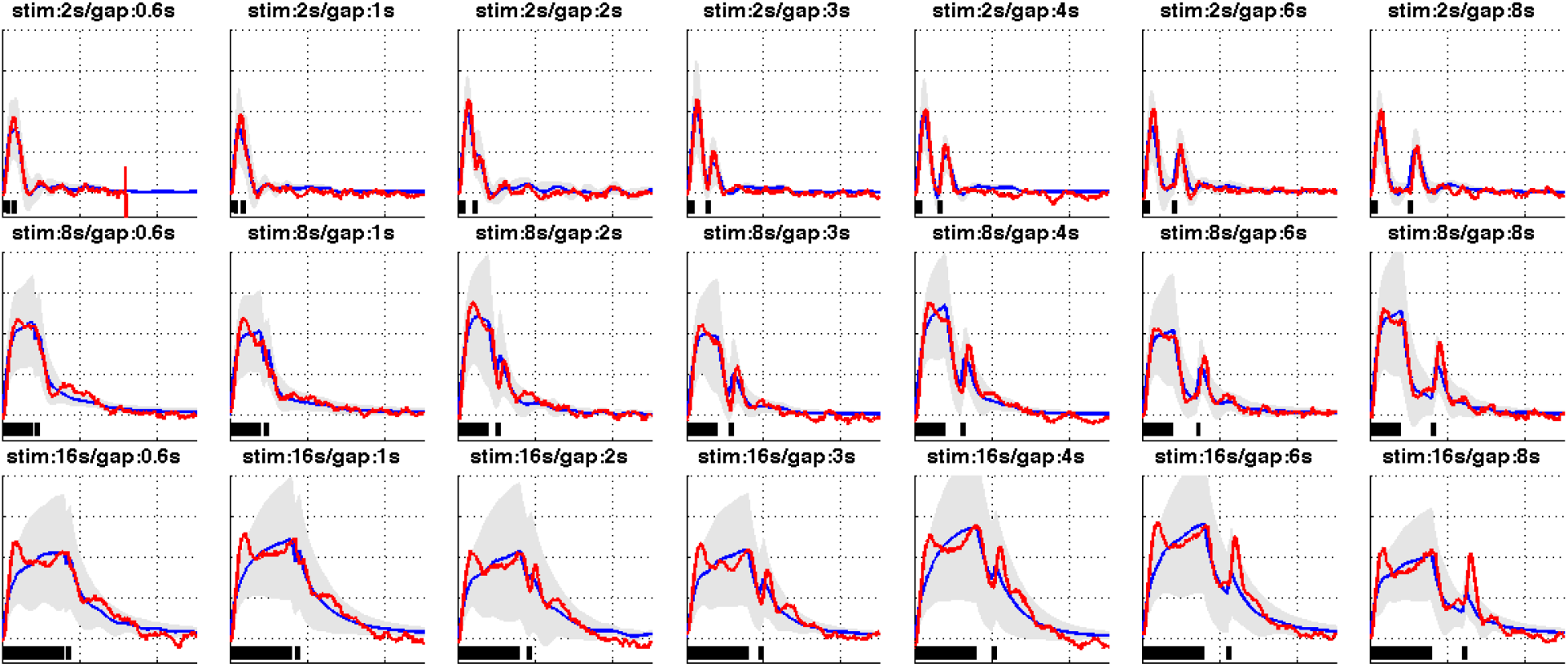
Average fit of the fractional-order model (shown in blue) to experimental CBF data (shown in red). The figures also illustrate the standard deviation of model estimates across all subjects (shown in gray). The lower bars indicate the stimulation time-courses. The figure shows a very good fit for the short stimulation paradigm (2s), a moderate fit for the 8s stimulation paradigm and a poorer fit for the long stimulation (16s) paradigm.

## IV. Discussion

In this investigation, we have shown that the fractional-order framework has a great potential in describing and characterizing a wide range of cerebral hemodynamic responses than the standard integer-order model. The fractional-order model’s ability to cover different response shapes is due to its flexibility and compliance offered by the two extra fractional order parameters, namely the fractional differentiation orders. Fractional parameters seem to characterize the CBF response in distinct ways by controlling the overshoots and the undershoots observed in the real CBF signal. Our sensitivity analysis clearly shows that these two parameters have different contributions in characterizing the cerebral hemodynamic determinants. While the variation of the positive overshoot of the CBF signal is sensitive to the value of the fractional differentiation order *q*_1_, the negative part of the CBF signal, namely the signal decay and undershoot, are more sensitive to the value of the second fractional differentiation order *q*_2_. In this study, we noticed that the model fails to fit the longer stimulation paradigm. When the framework involves highly non-linear dynamics, this fact may limit the model’s applicability to the long block design experiments of blocks longer than 8 seconds.

It is worth mentioning the strong correlation between *k_s_* and both *q*_1_ and *q*_2_ (see Fig. 3C and 3E). These correlations indicate that, within a specific interval, *k_s_* parameter can ”undo” the effect of both fractional-order parameters. This collinearity translates into a difficulty of accurately estimating *k_s_* in the fractional-order model.

Physiological interpretations of the fractional-order parameters *q*_1_ and *q*_2_ may be challenging. Similar to other NVC models, the model is meant to be descriptive to fit a wider range of experimental data that previous models cannot account for (see [4] for a discussion). Although these descriptive models have less physiologically interpretable parameters, they are extremely useful for comparison between different groups and conditions.

## V. Conclusion

Modeling the NVC mechanism is undertaking considerable development. It is essential for a better assessment of BOLD fMRI data and also a necessary step towards understanding the physiology behind the complex phenomena involved and finding biomarkers that represent the key features observed in measured CBF profiles. Current NVC models still lack the flexibility to fit a wider range of observed experimental data. In this paper, the framework of fractional calculus is used to model the CBF response to neural activity. A fractional-order oscillator is proposed based on the well-accepted and known minimal model proposed by Friston *et al*.. Through an optimization scheme that compares the original integer-order model and the proposed fractional one, we showed that the added fractional parameters provide a unique contribution in describing the CBF that can not be captured using the integer-order model’s parameters. Moreover, we assessed how sensitive CBF measure is to changes in the parameters of the model. Furthermore, using real neural activity-CBF data, the fractional model has proven capable of fitting wider CBF responses to both event and block design input paradigms. Although fractional model parameters are harder to interpret physiologically, they offer a great opportunity to compare groups or conditions.

This paper does not deal with the estimation problem of the model’s unknown parameters and fractional orders. This task may be the subject of a forthcoming paper where model-based estimation techniques for fractional systems will be proposed.

## Acknowledgement

The authors would like to thank Prof. Ying Zheng from the University of Sheffield, UK, who provided them with the real data that helped in conducting the work of Section II.

## Notes

### Competing Interest Statement

The authors have declared no competing interest.

